# Insula uses overlapping codes for emotion in self and others

**DOI:** 10.1101/2024.06.04.596966

**Authors:** Jiayang Xiao, Joshua A. Adkinson, Anusha B. Allawala, Garrett Banks, Eleonora Bartoli, Xiaoxu Fan, Madaline Mocchi, Bailey Pascuzzi, Suhruthaa Pulapaka, Melissa C. Franch, Sanjay J. Mathew, Raissa K. Mathura, John Myers, Victoria Pirtle, Nicole R Provenza, Ben Shofty, Andrew J. Watrous, Xaq Pitkow, Wayne K. Goodman, Nader Pouratian, Sameer Sheth, Kelly R. Bijanki, Benjamin Y. Hayden

## Abstract

In daily life, we must recognize others’ emotions so we can respond appropriately. This ability may rely, at least in part, on neural responses similar to those associated with our own emotions. We hypothesized that the insula, a cortical region near the junction of the temporal, parietal, and frontal lobes, may play a key role in this process. We recorded local field potential (LFP) activity in human neurosurgical patients performing two tasks, one focused on identifying their own emotional response and one on identifying facial emotional responses in others. We found matching patterns of gamma- and high-gamma band activity for the two tasks in the insula. Three other regions (MTL, ACC, and OFC) clearly encoded both self- and other-emotions, but used orthogonal activity patterns to do so. These results support the hypothesis that the insula plays a particularly important role in mediating between experienced vs. observed emotions.

## INTRODUCTION

The ability to understand and respond appropriately to emotions of other people is one of the key skills in humans’ cognitive repertoires (Happé et al., 2017; Preckel et al., 2018; Schurz et al., 2021). This ability in turn is central to social skills like empathy and theory of mind. Emotion identification is of great interest to psychiatrists because it is compromised in diseases like depression and autism (Fletcher-Watson & Bird, 2020; Gambin & Sharp, 2018; Nowacki et al., 2020). Consequently, the foundations of this skill have been of great interest in neuroscience and psychology (de Waal & Preston, 2017; Decety, 2015; Marsh, 2018). One important question is how recognizing our own emotions relates to recognizing emotions in others.

One possibility is that these two abilities rely on shared neural representations that can translate between representations of emotion in self and others. This idea is supported by findings from related fields. For example, the ability to identify visceral states in others depends in part on circuitry related to perceiving our own emotions (Preckel et al., 2018; Singer & Lamm, 2009; Singer & Tusche, 2014). When people experience a painful stimulus, part of the pain network including bilateral anterior insula and anterior cingulate cortex is activated as when they see a loved one have a similar experience (Lamm et al., 2011; Singer et al., 2004). Likewise, multivoxel pattern analysis found that feelings of disgust and perceptions of unfairness also co-activate the insula and cingulate cortex in similar ways (Corradi-Dell’Acqua et al., 2016). The overlapping activation is consistent with the hypothesis that observers infer the emotional states of others through a process that makes use of overlapping representations that help map the experiences of the observed onto the observers’ own state (de Waal & Preston, 2017). Indeed, some authors have conjectured that shared reactivations may be a core element of theory of mind and may contribute to empathy as well (Mitchell & Phillips, 2015; Molenberghs et al., 2016; Schaafsma et al., 2015).

Emotion activates much of the brain, including regions like the medial temporal lobe (MTL), anterior cingulate cortex (ACC), orbitofrontal cortex (OFC), and insula (Hultman et al., 2016; Janak & Tye, 2015; Rajmohan & Mohandas, 2007; Underwood et al., 2021). Among these regions, the insula is of particular interest (Gasquoine, 2014; Menon & Uddin, 2010). The insula responds to viscerally relevant stimuli, such as pain or disgust-inducing odor, whether experienced personally or observed in others (Kanske et al., 2015; Lamm et al., 2011; Singer et al., 2009; Zhao et al., 2022). The insula is a key node in the salience network (Menon & Uddin, 2010; Uddin, 2017), which shapes our emotional experiences and contributes to the generation of emotional states (Cisler et al., 2019; Luo et al., 2014; Underwood et al., 2021). Moreover, insula contributes to the perception of bodily states, which in turn are thought to be foundational to emotion processing (Craig, 2002; Karnath et al., 2005; Paulus & Khalsa, 2021). Indeed, activity in the insula is elicited by a variety of emotional events, suggesting a role beyond visceral sensory processing (Nguyen et al., 2016; Uddin, 2015; Zhou et al., 2021). However, limited accessibility and the rarity of isolated lesions have provided difficult barriers to a more sophisticated understanding of its role (Molnar-Szakacs & Uddin, 2022; Uddin, 2017).

We recorded local field potentials in the human insula, and, for comparison, the MTL, ACC, and OFC. Participants performed two tasks, an emotional identification task (in which they identified their own emotional response) and an emotional expression identification task (in which they identified another person’s emotional response). In the insula, we found a response pattern that encoded emotions and was consistent across the two tasks, highlighting the convergence and generalizability of valence representation. We found clear encoding of emotion in both tasks in the other three regions, but no evidence for overlapping representations of self- and other-emotions. Thus, within our set of four sampled regions, overlapping representations appear to be unique to the insula. Together, these findings suggest that the insula can process emotional valence across different person contexts and suggest a possible basis for empathetic cognition.

## RESULTS

### Task and recording sites

We recorded from intracranial electrodes while human epilepsy patients (n=17, nine female) viewed videos (*Emotional brain behavior quantification task*) and facial expressions (*Affective Bias task*) that varied along the emotional valence dimension (**Figure 1A**). The emotional brain behavior quantification (EBBQ) task was used to test for self-emotion and the Affective Bias task was used to test for other-emotion.

**Figure 1:**
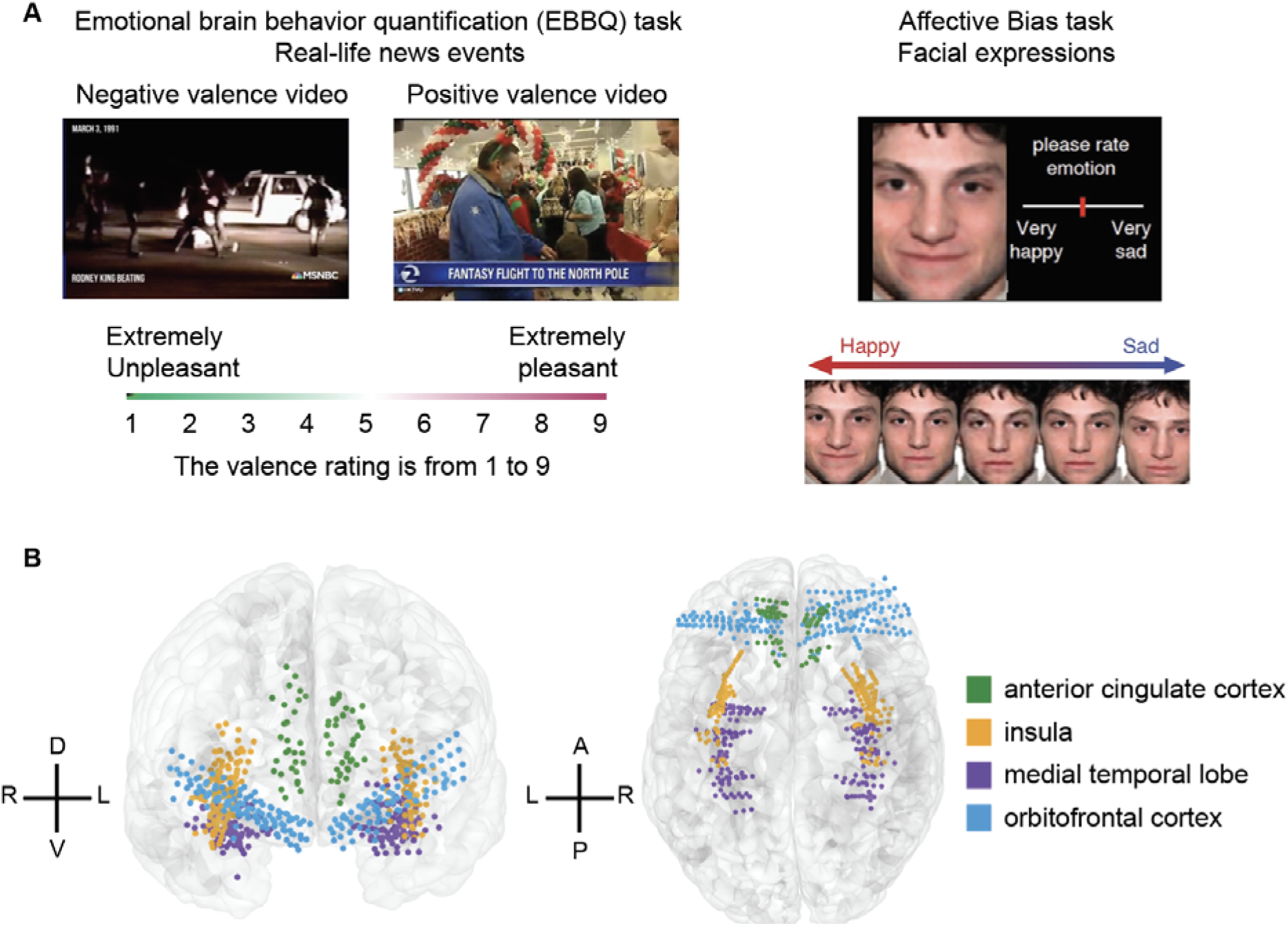
Self and other emotion tasks and neural recording locations. **(A)** Task structure. Each participant completed both EBBQ task (left) and Affective Bias task (right). **(B)** Recording locations across all participants on a template brain. Coordinates are in Montreal Neurological Institute (MNI) space. Different colors indicate contacts from different regions: green, anterior cingulate cortex; orange, insula; purple, medial temporal lobe; blue, orbitofrontal cortex. Dorsal (D), ventral (V), left (L), right (R), anterior (A), and posterior (P) directions are indicated.

In the EBBQ task, stimuli consisted of 30-60 second duration emotional videos containing real-life news events (Samide et al., 2020). We used real-world news clips because they are clearly non-fictional, and thereby provide greater validity than previous studies that used clips taken from movies or music videos (Abraham et al., 2008). Participants rated their emotional valence at the end of each video.

In the Affective Bias task, participants viewed a series of emotional facial expressions and rated the emotion of the face using a continuous scale by clicking on a slider bar (Bijanki et al., 2019; Metzger et al., 2023; Surguladze et al., 2004). The stimulus set consisted of computer-morphed faces spanning different emotional valence and intensity (100% happy, 50% happy, 30% happy, 10% happy, neutral, 10% sad, 30% sad, 50% sad, and 100% sad). Previous behavioral studies have shown that in this task, depression patients required a higher intensity of happiness to identify a face as happy compared to the healthy control group, and this perceptual bias was reduced with clinical improvement (Münkler et al., 2015).

### Response to emotional valence is heterogeneous across contacts

We recorded from 678 contacts in four brain regions in 17 participants who performed both tasks (**Figure 1B**). We first computed spectral power from each of six frequency bands: delta (1-4 Hz), theta (4–8 Hz), alpha (8–12 Hz), beta (12–30 Hz), gamma (35–50 Hz), and high-gamma (70–150 Hz) (These are standard definitions of these bands; we used the same band definitions in Xiao et al., 2023). For each band, we fit a linear regression model between spectral power and valence rating, using the least-squares approach, to obtain a normalized regression weight on the single contact level (**Figure 2A**). A positive normalized regression weight (that is, normalized beta value, shown in red in **Figure 2B-E**) indicates the contact has a larger response in gamma band during positive valence stimuli in each of the four regions we recorded. These illustrative figures highlight the diversity of responses across contacts within each region.

**Figure 2.**
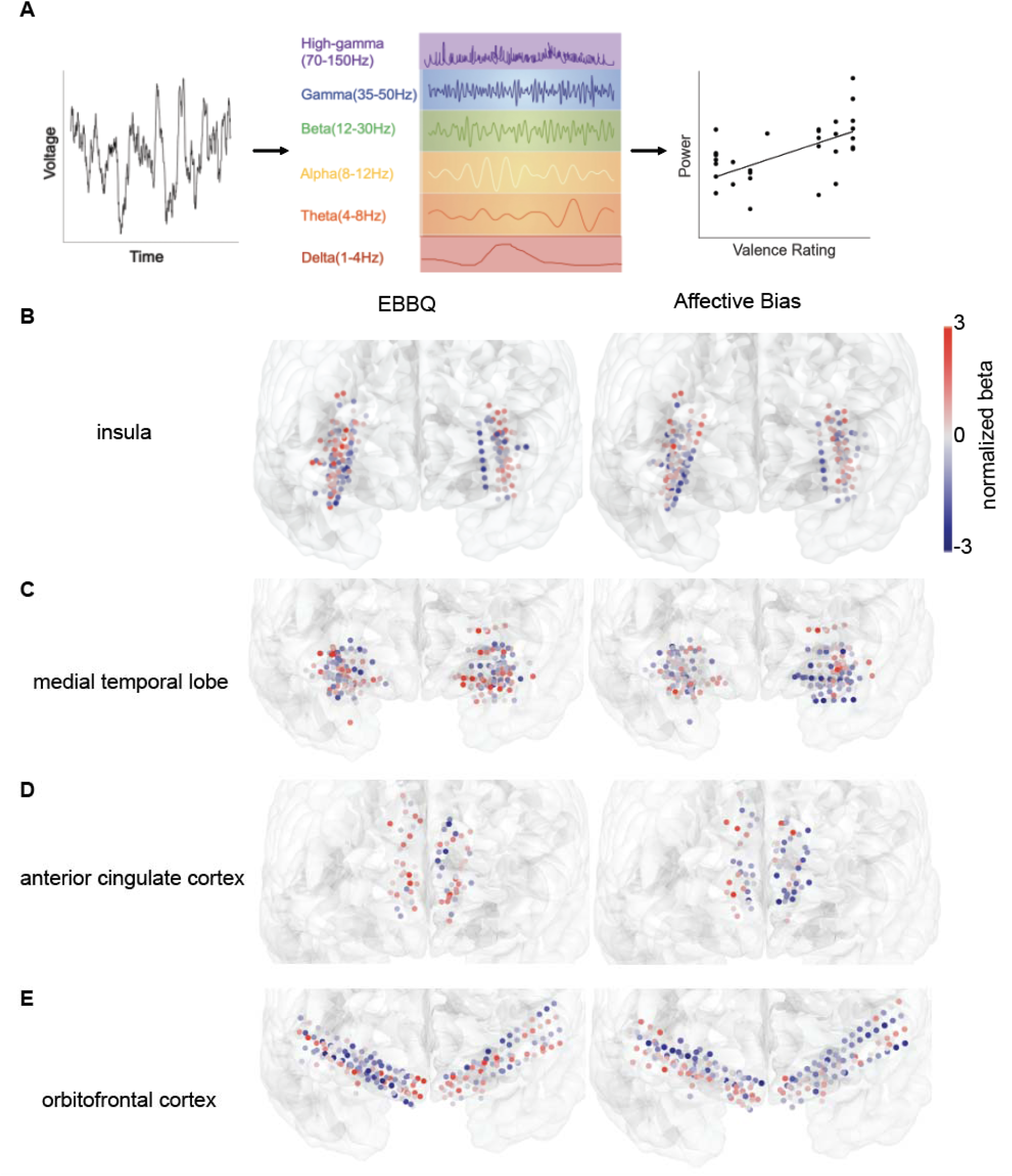
Neural response pattern across tasks. **(A)** For each contact, we obtained one normalized beta value for each frequency band representing the relationship between the spectral power in that band and the valence rating. **(B-E)** Normalized beta value for gamma power across contacts during EBBQ and Affective Bias task in insula **(B)**, medial temporal lobe **(C)**, anterior cingulate cortex **(D)**, and orbitofrontal cortex **(E)**. Contacts in red have a larger response during positive valence stimuli while contacts in blue have a larger response during negative valence stimuli.

We were especially interested in the gamma and high-gamma bands. Spectral power in these bands is a good index of local processing (Daume et al., 2024; Henrie & Shapley, 2005; Nir et al., 2007). Prior research has shown a close relationship between the spiking activity of neurons and the high-frequency activity observed in intracranial recordings (Lachaux et al., 2012; Nir et al., 2007; Rich & Wallis, 2017). The primary contributor to the high-frequency components of the local field potential may be the synchronized spiking of neighboring neurons (Buzsáki et al., 2012). Thus, it has been proposed that spectral power in high-frequency bands including gamma and high-gamma may represent the collective activity of a local group of neurons, and could potentially serve as a potential proxy for neuronal activity (Colgin et al., 2009; Fernández-Ruiz et al., 2021; Manning et al., 2009; Miller et al., 2014).

In the insula, we found clear encoding of emotional valence in both tasks. Specifically, the spectral power in gamma or high-gamma band was modulated by emotional valence in 13.5% of contacts during the EBBQ task, and 11.1% was modulated during the Affective Bias Task (EBBQ: n=28/208 contacts, Affective Bias Task: n=23/208 contacts, note that these proportions were generated using *p*-values Bonferroni corrected for two frequency bands). Both of these proportions were substantially greater than would be expected by chance (chance level: 5.0%, *p* < 0.05 when compared to the true proportions).

Valence was also encoded in both tasks in the three other regions in our study: the medial temporal lobe (MTL, EBBQ: n=29/198, 14.6%, Affective Bias Task: n=23/198, 11.6%), the anterior cingulate cortex (ACC, EBBQ: n=11/69, 15.9%, Affective Bias Task: n=15/69, 21.7%), and the orbitofrontal cortex (OFC, EBBQ: n=27/203, 13.3%, Affective Bias Task: n=29/203, 14.3%). These proportions from all regions were significantly larger than expected by chance (chance level: 5.0%, *p* < 0.05 in all cases).

Overall, the proportion of contacts encoding emotional valence was similar between the EBBQ task and the Affective Bias Task. Indeed, we did not find any significant difference in the encoding of valence across tasks in all four regions (*p* > 0.9 for all four regions, chi-squared test). In both the EBBQ and the Affective Bias task, and in all regions, the spectral power was greater during positive valence in some contacts while in other contacts the power was greater during negative valence; in other words, the response is heterogeneous across contacts (**Figure 2B-E**). Collectively, these results indicate that emotion is encoded in all four regions studied, consistent with findings from earlier studies (Jackson et al., 2024; Rogers-Carter et al., 2018; Rolls, 2019; Zheng et al., 2019).

### Task-general and task-specific coding

We next asked whether the pattern of activation across contacts was similar within each brain region for the two tasks. To quantify the similarity between tasks, we calculated the correlation between the normalized regression weights (betas) from the EBBQ task and the normalized betas from the Affective Bias task. We concatenated the lists of regression weights into two vectors for each brain region, one for each task. Each element in a vector therefore corresponded to a specific recording site within a brain region. Correlating these two vectors essentially asks whether variation in one variable covaries with variation in the other; a positive correlation would suggest a *common code* for empathy between the two tasks (cf. Azab & Hayden, 2017). This analysis therefore assesses the similarity in the pattern of activation across the two conditions.

In the insula, we found a positive correlation between these normalized regression weights, indicating that the response pattern in high-frequency bands is consistent across tasks (**Figure 3A**, gamma: r = 0.34, *p* = 4.24×10^-6^; high-gamma: r = 0.32, *p* = 2.79×10^-5^). In the three other brain regions, we did not observe a significant positive correlation between normalized betas across tasks (**Figure 3B-D**, *p* > 0.9 for each of these regions). The measured correlation coefficients within the insula were substantially larger than those of other brain regions (Fisher’s z-transform, *p* values were Bonferroni corrected for gamma and high-gamma bands; compared with MTL: z = 4.19, *p* = 5.70×10^-5^ for gamma band, z = 2.59, *p* = 0.020 for high-gamma band; compared with ACC: z = 2.83, *p* = 0.0092 for gamma band, z = 3.04, *p* = 0.0047 for high-gamma band; compared with OFC: z = 4.15, *p* = 6.77×10^-5^ for gamma band, z = 3.67, *p* = 4.81×10^-4^ for high-gamma band).

**Figure 3.**
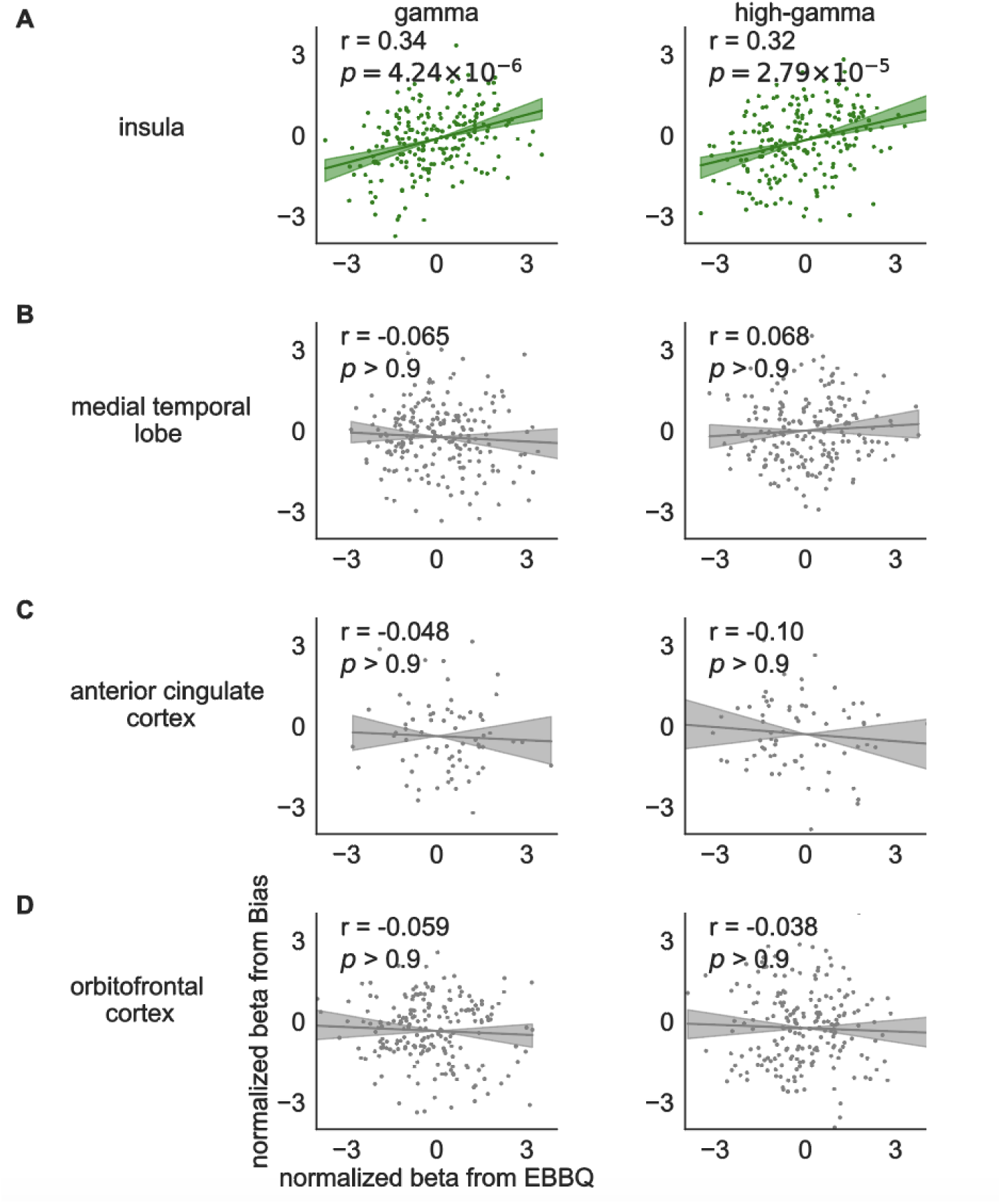
Cross-task coding in gamma and high-gamma activity. **(A)** Correlation between the normalized betas for gamma and high-gamma power in insula from EBBQ task and the normalized betas from Affective Bias task. Each dot indicates one contact. **(C-D)** Correlation between the normalized betas from EBBQ task and Affective Bias task in other brain regions. Regions with significant positive correlations were shown in green while non-significant results were shown in gray.

We next asked whether the positive correlation between these normalized betas in the insula reflects a combination of abstract and task-specific codes. To do so we used a positive-control / negative-control approach we have used in the past (Fine et al., 2023; Herman et al., 2023; Yoo and Hayden, 2020). We split the data within a single task into two halves and generated bootstrapped beta values from each half of the data. The correlation between these vectors provides an estimate of the correlation that would be measured given the noise properties of the stimuli under the assumption that the correlation was perfect (see also Azab & Hayden, 2017, 2018). We repeated the entire process 1000 times to generate a distribution of randomized perfectly correlated r-values. We then examined where the experimental r-value we obtained from the across-task correlation fell within this positive control distribution. In the medial temporal lobe, anterior cingulate cortex, and orbitofrontal cortex, the experimental r-values we obtained from the across-task correlation were significantly lower than r-values from the correlations within tasks (*p* < 0.05 in these brain regions). However, we did not observe a significant difference between these r-values in the insula (*p* > 0.5 in the insula). The insula result indicates that its encoding is, to the best of our ability to measure, abstract, and does not contain a clearly detectable task-selective component. Conversely, the other regions do not have a measurable task-general component.

In other words, these results endorse a potential qualitative difference between the insula and the other regions we studied, in which the insula has an abstract (i.e., cross-task) encoding of valence and the other regions do not. While the other regions encode emotion and do so with a similar magnitude to the insula, we see no evidence that their code is the same across the two contexts of first-person (self emotions) and second-person (others’ emotions). Thus, within our four regions, the insula appears the most likely substrate for cross-person emotional encoding.

### Valence is decodable both within task and across tasks in the human insula

We next used decoding analysis to test whether the valence representation is decodable across tasks in these regions. We performed both within-task and cross-task decoding to predict valence condition (positive/negative) using the average spectral power across the trial in all frequency bands (**Figure 4A**). For within-task analysis, we used training and testing data from the same task and performed stratified 10-fold cross-validation using a support vector classifier (Berrar, 2018; Chang & Lin, 2011). For cross-task generalization analysis, we used training and testing data from different tasks.

**Figure 4.**
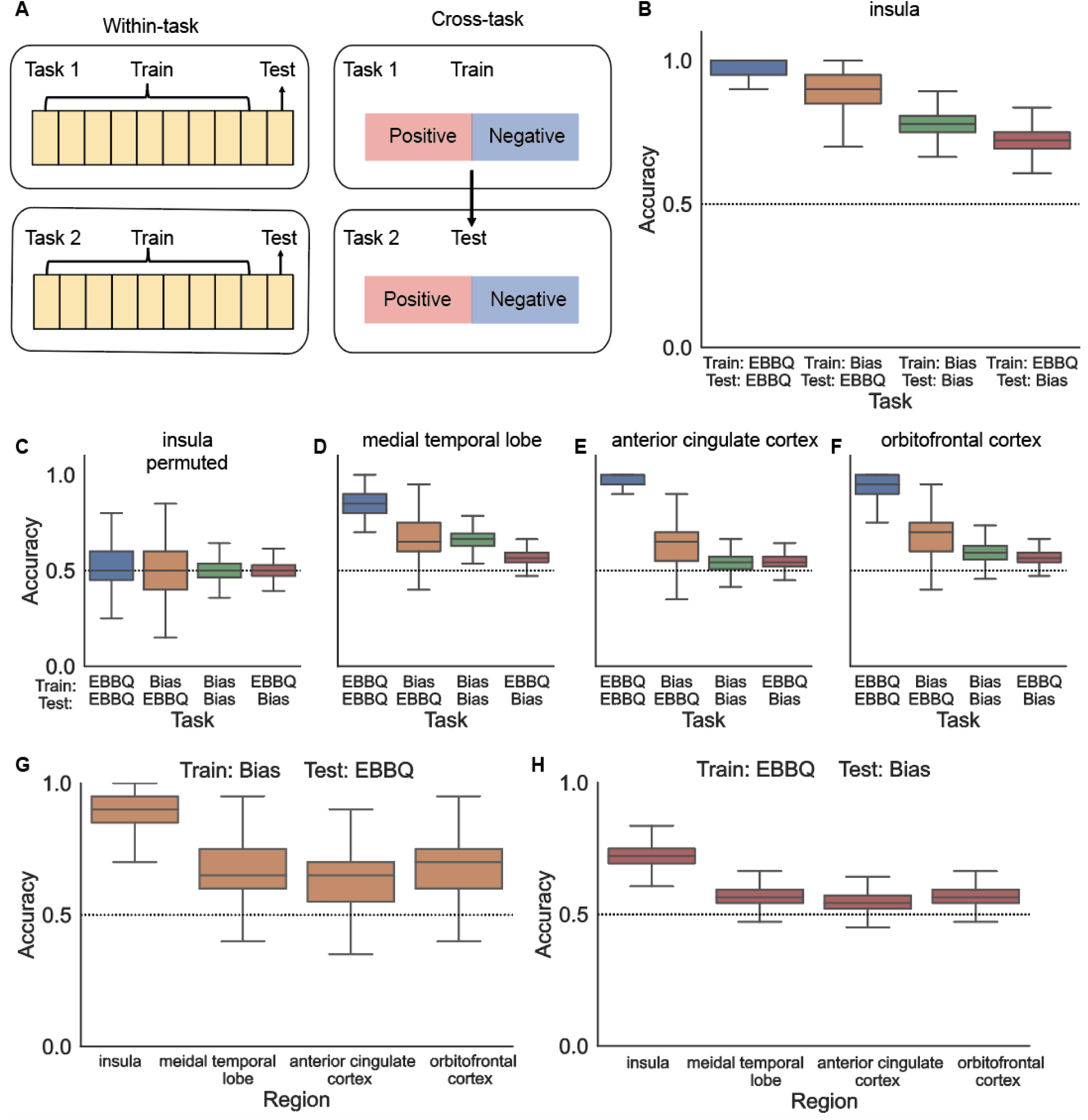
Decoding valence condition from neural activity. **(A)** Analysis paradigm. **(B)** Decoding accuracy for each training and testing pair using insula data. Boxplots illustrate quartiles at 25% and 75%, with horizontal lines denoting medians, and whiskers extending to 1.5 times the interquartile ranges. The dotted line indicates the chance level performance (50% accuracy). **(C)** Decoding accuracy using permuted insula data. **(D-F)** Decoding accuracy using data in medial temporal lobe, anterior cingulate cortex, and orbitofrontal cortex. **(G-H)** Comparison of decoding accuracy across regions for the same type of training and testing pairs.

First, using only the insula data, we found that accuracies for all training and testing pairs were higher than chance (**Figure 4B**, *p* < 0.001 for all pairs after Bonferroni correction for four regions when compared with an accuracy of 0.50). This was true in both tasks separately. Interestingly, EBBQ was overall more decodable than bias; however, both EBBQ and Bias were highly significant. To confirm that our findings were not artifactual, we permuted the valence condition labels and repeated the same decoding process 10000 times. With the valence ratings permuted and thus mismatched with the neural data, the valence condition could no longer be accurately predicted (**Figure 4C**, *p* > 0.5 for all pairs when compared with an accuracy of 0.50). In any case, the highly significant cross-task decoding indicates that the valence condition can be decoded by the decoder trained on a different task, consistent with a pattern of abstract valence encoding in the insula.

In comparison, we performed the decoding analysis for the other three brain regions: medial temporal lobe (**Figure 4D**, *p* values from left to right when compared with an accuracy of 0.50, Bonferroni corrected for four regions: 0.003, 0.36, 0.003, 0.24), anterior cingulate cortex (**Figure 4E**, *p* values from left to right: <0.001, 0.80, 0.91, 0.47), orbitofrontal cortex (**Figure 4F**, *p* values from left to right: <0.001, 0.32, 0.17, 0.20). Most notably, the cross-task decoding was not statistically significant for either direction in any of the three brain regions. We performed Kruskal-Wallis tests to assess significant differences in decoding accuracy across regions (**Figure 4G-H**, train Bias, test EBBQ: H = 20070.01, *p* < 1×10^-4^; train EBBQ, test Bias: H = 23319.37, *p* < 1×10^-4^). Subsequent post-hoc Wilcoxon rank sum tests confirmed that the accuracies for training and testing pairs from different tasks, using data from the insula, were much higher than the accuracies obtained from other brain regions (Bonferroni-corrected, *p* < 1×10^-4^ for insula vs. MTL, insula vs. ACC, insula vs. OFC). The higher accuracy in the insula indicates that the abstract representation is stronger in the insula than in other regions.

## DISCUSSION

We recorded intracranial field potentials while human neurosurgical patients performed a task in which they responded to emotional videos and, in a separate task, evaluated facial expressions for emotion. We found that in the insula, much more than other regions tested (MTL, ACC, and OFC), neural activity was similar for similar emotions in the two tasks, and thus for self and other. Thus, our results suggest that the insula carries a cross-person representation of valence, thereby expanding our understanding of its role in emotional processing. This type of cross-task commonality suggests that the insula carries an abstract encoding of emotional state, independent from the person it is associated with. This abstraction may help us in drawing conclusions about the emotional states of others (de Waal & Preston, 2017; Preckel et al., 2018).

These results are related to questions about how we recognize emotions in others and how the ability to do so relates to our own emotional states (Ferretti & Papaleo, 2019; Spunt & Adolphs, 2019). Notably, in simulation-based theories, we can recognize emotional states in other people by simulating the expression of emotions, that is, by recapitulating them (Goldie, 1999; Marsh, 2018). If this theory is correct, then it would help quite a bit to have a part of the brain in which we have an abstract representation of the emotion, so that it can be applied from self to other. Our results therefore suggest that the insula may be one such site in the brain.

The role of the insula in emotional processing remains a subject of ongoing debate. The insula is activated by a wide range of emotional stimuli, from acute painful sensations to fearful facial expressions (Frot et al., 2022; Segerdahl et al., 2015). However, some researchers have proposed that the insula primarily serves as a sensory rather than affective region, with other regions such as the orbitofrontal cortex playing a larger role in value representation in subsequent stages of processing (Rolls, 2016). A previous fMRI study, for instance, found that the anterior insula specifically encodes valence related to taste, but not vision, whereas the orbitofrontal cortex encodes valence independent of the sensory source (Chikazoe et al., 2014). Our results challenge this idea.

Our results do not, of course, demonstrate that the other regions from which we recorded have no role in emotional processing. Indeed, our results highlight the broadly distributed nature of emotional information throughout the four regions from which we recorded. We suspect that emotion, which has a pervasive influence on our decisions and our interpretation of the world, is likely represented broadly throughout the brain. If so, it would align with reward, which is closely related to emotion, and is likewise represented in many brain regions simultaneously (Fine & Hayden, 2021; Vickery et al., 2011). Indeed, we believe that the demonstration of both self and other emotion encoding in these regions argues against a strong functional specialization for different aspects of emotion (Westlin et al., 2023).

To make our stimuli more relevant to real life, and thereby increase the ecological validity of our findings, we used videos containing non-fictional news clips as part of the study’s stimuli (Samide et al., 2020). However, the use of emotional videos and facial expressions may not fully capture the intricate nature of emotional processing. Emotion is a multifaceted phenomenon that involves various sensory modalities, including visual, auditory, tactile, and olfactory components (Schirmer & Adolphs, 2017; Feldman Barrett, 2006). Considering the involvement of the insula in gustatory and olfactory processing, as well as its role in multisensory integration, future research should consider incorporating additional sensory modalities to provide a more comprehensive understanding of how the insula processes emotions (Chikazoe et al., 2019; Gogolla, 2017; Koeppel et al., 2020).

The perception of emotional stimuli not only enriches our daily experiences but also plays a vital role in our survival. Emotional valence processing could influence perceptual, attentional, and mnemonic processes, thus guiding our behaviors and shaping our interactions with the world around us (Ballhausen et al., 2015; Dolcos et al., 2020; Gutchess & Kensinger, 2018). Disruptions in this crucial process have been strongly associated with various psychiatric conditions (Price & Duman, 2020; Tschida & Yerys, 2021). For example, neutral or ambiguous facial expressions tend to be evaluated as sad in individuals with depression (Bourke et al., 2010; Disner et al., 2011). This alternation in valence processing can profoundly impact how they perceive and interact with their environment. In the future, researchers can explore potential changes in neural activity among these patients. Understanding the neural underpinnings of emotional valence processing in healthy participants and individuals with psychiatric disorders may provide valuable insights for early detection and targeted treatment, ultimately offering important clinical benefits. Ultimately, our results provide a valuable complement to studies using animal models and studies using non-invasive methods in humans. These kinds of studies have had a great impact, but for studying certain high-level aspects of cognition, it is valuable to have human participants and direct brain measures (Widge et al., 2019).

## EXPERIMENTAL METHODS

### Participants

Seventeen participants (eight males and nine females, mean age 43 years, range 19-63 years) undergoing invasive monitoring for the treatment of refractory epilepsy at Baylor St. Luke’s Medical Center (Houston, Texas, USA) participated in our study. The implantation sites were exclusively determined by the clinical team with the sole goal of localizing the seizure onset zone. Ethical approval for the study was granted by the Institutional Review Board at Baylor College of Medicine, under IRB protocol number H-18112. All participants provided both verbal and written consent to participate in the study. After the surgical implantation of electrodes, patients underwent approximately one week of inpatient monitoring. During this period, we conducted both the EBBQ task and the Affective Bias (in the same patients) task while concurrently recording neural activity. These tasks were presented on a Viewsonic VP150 monitor with a resolution of 1920 × 1080, positioned 57 cm away from the participants.

### Emotional brain behavior quantification (EBBQ) task

Each run in this task contained eight emotionally positive and eight emotionally negative videos. The valence category was predefined by average ratings from 100 healthy students in a publicly available database and all videos contained real-life news events (Samide et al., 2020). Unlike previous studies that have used video clips from movies or music videos, which might be processed differently due to their fictional nature, the video database employed in our research used real-life newscasts. At the end of each video, participants provided a valence rating using a USB numeric keypad, ranging from 1 (extremely unpleasant) to 9 (extremely pleasant). In this dataset, 16 out of the 17 participants completed two runs of this task, while the remaining participant completed four runs. Data from all runs were included in the analysis.

### Affective Bias task

Participants were asked to rate emotional human face photographs. We used a set of happy, sad, and neutral face examples (6 identities for each emotion, split evenly between genders), adapted from the NimStim Face Stimulus Set (Tottenham et al., 2009). These faces were manipulated through a Delaunay tessellation matrix to create subtle variations in emotional intensity, ranging from neutral to highly expressive. The increments were set at 10%, 30%, 50%, and 100% for happy and sad expressions respectively.

In each trial, a white fixation cross was presented on a black background for 1000 msec (jittered +/− 100 msec). Then a face image and a rating prompt were displayed simultaneously on the screen. The rating prompt included an interactive analog slider bar positioned under the text instruction: “Please rate the emotion”. Participants indicated their rating by clicking a specific position on a slider bar using a computer mouse. The ratings were captured on a continuous scale ranging from 0 (’Very Sad’) to 0.5 (’Neutral’) to 1 (’Very Happy’). The presentation of stimuli followed a blocked design, with all happy faces (plus neutral) appearing in one block while all sad faces (plus neutral) appearing in a separate block. Each run consisted of one block of happy faces and one block of sad faces, with each block containing 30 randomized trials (6 identities x 5 levels of intensity). Participants completed between two to twelve runs with alternating happy and sad trials. All data from these runs were included in the analysis. The order of these blocks, either starting with happy faces or sad faces, was counterbalanced across participants.

### Electrode localization and intracranial recordings

Prior to surgery, patients underwent brain magnetic resonance imaging (MRI), followed by post-implantation computed tomography (CT). For precise localization of implanted electrodes, we aligned the pre-operative T1-weighted MRI scans with the post-operative CT scans. The automated cortical reconstruction was executed using Freesurfer tools (Fischl, 2012). After mapping the electrodes onto the native MRI space using BioImage Suite v3.5b1 (Joshi et al., 2011), an expert anatomist assessed the images to identify brain regions and categorized contacts as either gray or white matter. Contacts found within white matter were excluded from subsequent analyses. Detailed methodologies have been outlined elsewhere (Sheth et al., 2022; Xiao et al., 2023). Neural signals were recorded using sEEG electrodes at a sampling rate of 2000 Hz through the Cerebus data acquisition system (BlackRock Microsystems, UT, USA). The signals were processed with a bandpass filter set at 0.3-500 Hz using a 4th-order Butterworth filter.

### Preprocessing and spectral decomposition

We conducted a visual inspection of the raw signals to identify any recording artifacts and interictal epileptic spikes. Contacts exhibiting excessive noise were excluded to prevent noise propagation to other contacts through re-referencing. To minimize the impact of line-noise artifacts and volume conduction, signals were notch-filtered (60 Hz and its harmonics) and then re-referenced through bipolar referencing (Bastos & Schoffelen, 2015). Subsequently, the referenced signals were down-sampled to 1000 Hz, followed by the application of a Hilbert transform to estimate spectral power across six distinct frequency bands: 1-4 Hz (delta), 4–8 Hz (theta), 8–12 Hz (alpha), 12–30 Hz (beta), 35–50 Hz (gamma), and 70–150 Hz (high-gamma).

### Calculation of spectral power

To determine spectral power values for each trial, we computed the average of squared magnitudes from the Hilbert transform decomposition. For the EBBQ task, spectral power was averaged across a time window beginning at video onset and ending at video offset. For the Affective Bias task, the mean spectral power was calculated across a time window beginning at the onset of the face stimulus and ending at the participant’s response. The spectral power values were then standardized across trials in the same run for each contact.

### Data visualization in the template brain

Contacts were mapped onto the MNI space and visualized using the open-source software RAVE (R Analysis and Visualization of iEEG) (Magnotti et al., 2020). To assess variations in neural responses to stimuli of different valence ratings, we used an ordinary least squares regression to generate one normalized regression weight (normalized beta) for each contact. These normalized betas, which compared the average spectral power during positive valence videos/faces with that during negative valence videos/faces, were then plotted at each contact.

### Encoding of emotional valence in each task

A contact was considered selective for emotional valence when the spectral power in at least one frequency band was significantly modulated by valence in the linear regression model. To assess whether the proportion of selective contacts was significant, we derived a null distribution by repeating the same procedures after randomly permuting the valence ratings 1000 times. The p-value was defined as the probability that the proportion of selective contacts from the null distribution is higher than the true proportion. To test the encoding of emotional valence in both tasks, this analysis was conducted separately for each task in all four regions.

### Calculation of difference in correlation coefficient between regions

To compare the correlation coefficients, we transformed the correlation coefficients to Fisher’s z scores using the following equation for each correlation coefficient:

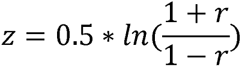

Then we computed the Z-test statistic for the difference between the two z scores using the following equation:

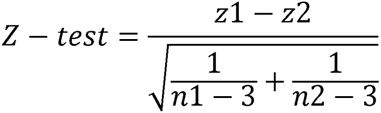

where z_1_ and z_2_ are the z scores transformed from the correlation coefficients for the two regions, and n_1_ and n_2_ are the sample sizes associated with the two values (number of contacts in each region).

Lastly, we determined whether the difference between the two correlation coefficients is statistically significant by comparing the Z-test statistic to the critical value from the standard normal distribution. A *p*-value of less than 0.05 indicates that the two correlation coefficients are significantly different from each other.

### Decoding

First, we identified all contacts within a specific brain region that had recordings from a minimum of ten trials for both positive and negative valence conditions. From this pool of available contacts, a subset of 100 contacts was randomly chosen. Due to the limited number of contacts in the anterior cingulate cortex, 50 contacts instead of 100 contacts were randomly chosen for the decoding analysis in this region. From each of these selected contacts, we randomly extracted spectral power values from ten EBBQ trials associated with either positive or negative valence conditions. By concatenating the spectral power of these 100 contacts across each of the ten trials, we generated a total of 10 * 2 data points. For the Affective Bias task, we randomly extracted spectral power values from 70 trials associated with either positive or negative valence conditions to generate a total of 70 * 2 data points.

To evaluate the within-task classification accuracy, we employed a stratified 10-fold cross-validation approach, ensuring that the distribution of samples for each valence condition was preserved in each fold. We employed the sklearn.svm.SVC class in Python’s Scikit-learn library to perform classification tasks using a support vector machine with a linear kernel. Predictions were generated using the trained classifier on the testing dataset through the ‘predict’ method.

Cross-task decoding is used to investigate whether the information encoded in the brain’s activity patterns during one task can be leveraged to accurately classify the brain’s responses during a different task. We trained the model using the same set of data from either EBBQ task or Affective Bias task and used the trained model to predict the valence label in the other task.

We then replicated this entire procedure 10000 times to attain a robust and reliable classification accuracy estimate. We counted the number of samples in the distribution for which the accuracy due to chance was higher than the true accuracy. The *p*-value was defined as the probability that the prediction accuracy is higher than 0.5.

We used a permutation test to further validate our model. By permuting the valence condition labels to mismatch it with the neural data and repeating the same decoding process 10000 times, we tested whether the valence condition could still be accurately predicted.

## Acknowledgments

We thank our patients for their participation in the research. This work was supported by the National Institutes of Health (Grant No. UH3 NS103549 [to SAS, KRB, WKG, and NP], Grant No. K01 MH116364 [to KRB], Grant No. R01 MH127006 [to KRB], Grant No. UH3 NS100549 [to WKG and SAS], and Grant No. R01 MH114854 [to WKG]) and McNair Foundation (to SAS).

## Author contributions

Conceptualization, JX, SAS, KRB, and BYH; methodology, JX, JAA, and BYH; investigation, JX, EB, XF, MM, BP, SP, and AJW; visualization, JX, GB, BP, SP, RKM, and BS; analysis, JX; writing – original draft, JX; writing – review & editing, all authors; funding acquisition, WKG, NP, SAS, and KRB; supervision, SAS, KRB, and BYH

## References

Abraham, A., von Cramon, D. Y., & Schubotz, R. I. (2008). Meeting George Bush versus meeting Cinderella: The neural response when telling apart what is real from what is fictional in the context of our reality. Journal of Cognitive Neuroscience, 20(6), 965–976. 10.1162/jocn.2008.20059

Azab, H., & Hayden, B. Y. (2017). Correlates of decisional dynamics in the dorsal anterior cingulate cortex. PLOS Biology, 15(11), e2003091. 10.1371/journal.pbio.2003091

Azab, H., & Hayden, B. Y. (2018). Correlates of economic decisions in the dorsal and subgenual anterior cingulate cortices. The European Journal of Neuroscience, 47(8), 979–993. 10.1111/ejn.13865

Ballhausen, N., Rendell, P. G., Henry, J. D., Joeffry, S., & Kliegel, M. (2015). Emotional valence differentially affects encoding and retrieval of prospective memory in older adults. *Aging*, Neuropsychology, and Cognition, 22(5), 544–559. 10.1080/13825585.2014.1001316

Bastos, A. M., & Schoffelen, J.-M. (2015). A Tutorial Review of Functional Connectivity Analysis Methods and Their Interpretational Pitfalls. Frontiers in Systems Neuroscience, 9, 175. 10.3389/fnsys.2015.00175

Berrar, D. (2018). Cross-Validation. 10.1016/B978-0-12-809633-8.20349-X

Bijanki, K. R., Manns, J. R., Inman, C. S., Choi, K. S., Harati, S., Pedersen, N. P., Drane, D. L., Waters, A. C., Fasano, R. E., Mayberg, H. S., & Willie, J. T. (2019). Cingulum stimulation enhances positive affect and anxiolysis to facilitate awake craniotomy. The Journal of Clinical Investigation, 129(3), 1152–1166. 10.1172/JCI120110

Bourke, C., Douglas, K., & Porter, R. (2010). Processing of facial emotion expression in major depression: A review. The Australian and New Zealand Journal of Psychiatry, 44(8), 681–696. 10.3109/00048674.2010.496359

Buzsáki, G., Anastassiou, C. A., & Koch, C. (2012). The origin of extracellular fields and currents—EEG, ECoG, LFP and spikes. Nature Reviews Neuroscience, 13(6), Article 6. 10.1038/nrn3241

Chang, C.-C., & Lin, C.-J. (2011). LIBSVM: A library for support vector machines. ACM Transactions on Intelligent Systems and Technology, 2(3), 27:1-27:27. 10.1145/1961189.1961199

Chikazoe, J., Lee, D. H., Kriegeskorte, N., & Anderson, A. K. (2014). Population coding of affect across stimuli, modalities and individuals. Nature Neuroscience, 17(8), Article 8. 10.1038/nn.3749

Chikazoe, J., Lee, D. H., Kriegeskorte, N., & Anderson, A. K. (2019). Distinct representations of basic taste qualities in human gustatory cortex. Nature Communications, 10(1), Article 1. 10.1038/s41467-019-08857-z

Cisler, J. M., Esbensen, K., Sellnow, K., Ross, M., Weaver, S., Sartin-Tarm, A., Herringa, R. J., & Kilts, C. D. (2019). Differential Roles of the Salience Network During Prediction Error Encoding and Facial Emotion Processing Among Female Adolescent Assault Victims. Biological Psychiatry: Cognitive Neuroscience and Neuroimaging, 4(4), 371–380. 10.1016/j.bpsc.2018.08.014

Colgin, L. L., Denninger, T., Fyhn, M., Hafting, T., Bonnevie, T., Jensen, O., Moser, M.-B., & Moser, E. I. (2009). Frequency of gamma oscillations routes flow of information in the hippocampus. Nature, 462(7271), 353–357. 10.1038/nature08573

Corradi-Dell’Acqua, C., Tusche, A., Vuilleumier, P., & Singer, T. (2016). Cross-modal representations of first-hand and vicarious pain, disgust and fairness in insular and cingulate cortex. Nature Communications, 7(1), Article 1. 10.1038/ncomms10904

Craig, A. D. (2002). How do you feel? Interoception: the sense of the physiological condition of the body. Nature Reviews Neuroscience, 3(8), Article 8. 10.1038/nrn894

Daume, J., Kamiński, J., Schjetnan, A. G. P., Salimpour, Y., Khan, U., Kyzar, M., Reed, C. M., Anderson, W. S., Valiante, T. A., Mamelak, A. N., & Rutishauser, U. (2024). Control of working memory by phase–amplitude coupling of human hippocampal neurons. Nature, 1–9. 10.1038/s41586-024-07309-z

de Waal, F. B. M., & Preston, S. D. (2017). Mammalian empathy: Behavioural manifestations and neural basis. Nature Reviews Neuroscience, 18(8), Article 8. 10.1038/nrn.2017.72

Decety, J. (2015). The neural pathways, development and functions of empathy. Current Opinion in Behavioral Sciences, 3, 1–6. 10.1016/j.cobeha.2014.12.001

Disner, S. G., Beevers, C. G., Haigh, E. A. P., & Beck, A. T. (2011). Neural mechanisms of the cognitive model of depression. Nature Reviews Neuroscience, 12(8), Article 8. 10.1038/nrn3027

Dolcos, F., Katsumi, Y., Moore, M., Berggren, N., de Gelder, B., Derakshan, N., Hamm, A. O., Koster, E. H. W., Ladouceur, C. D., Okon-Singer, H., Pegna, A. J., Richter, T., Schweizer, S., Van den Stock, J., Ventura-Bort, C., Weymar, M., & Dolcos, S. (2020). Neural correlates of emotion-attention interactions: From perception, learning, and memory to social cognition, individual differences, and training interventions. Neuroscience & Biobehavioral Reviews, 108, 559–601. 10.1016/j.neubiorev.2019.08.017

Feldman Barrett, L. (2006). Are emotions natural kinds?. Perspectives on psychological science, 1(1), 28–58.

Fernández-Ruiz, A., Oliva, A., Soula, M., Rocha-Almeida, F., Nagy, G. A., Martin-Vazquez, G., & Buzsáki, G. (2021). Gamma rhythm communication between entorhinal cortex and dentate gyrus neuronal assemblies. Science, 372(6537), eabf3119. 10.1126/science.abf3119

Ferretti, V., & Papaleo, F. (2019). Understanding others: Emotion recognition in humans and other animals. *Genes*, Brain and Behavior, 18(1), e12544. 10.1111/gbb.12544

Fine, J. M., & Hayden, B. Y. (2021). The whole prefrontal cortex is premotor cortex. Philosophical Transactions of the Royal Society B: Biological Sciences, 377(1844), 20200524. 10.1098/rstb.2020.0524

Fine, J. M., Maisson, D. J.-N., Yoo, S. B. M., Cash-Padgett, T. V., Wang, M. Z., Zimmermann, J., & Hayden, B. Y. (2023). Abstract Value Encoding in Neural Populations But Not Single Neurons. The Journal of Neuroscience: The Official Journal of the Society for Neuroscience, 43(25), 4650–4663. 10.1523/JNEUROSCI.1954-22.2023

Fischl, B. (2012). FreeSurfer. NeuroImage, 62(2), 774–781. 10.1016/j.neuroimage.2012.01.021

Fletcher-Watson, S., & Bird, G. (2020). Autism and empathy: What are the real links? Autism, 24(1), 3–6. 10.1177/1362361319883506

Frot, M., Mauguière, F., & Garcia-Larrea, L. (2022). Insular dichotomy in the implicit detection of emotions in human faces. Cerebral Cortex, 32(19), 4215–4228. 10.1093/cercor/bhab477

Gambin, M., & Sharp, C. (2018). The relations between empathy, guilt, shame and depression in inpatient adolescents. Journal of Affective Disorders, 241, 381–387. 10.1016/j.jad.2018.08.068

Gasquoine, P. G. (2014). Contributions of the Insula to Cognition and Emotion. Neuropsychology Review, 24(2), 77–87. 10.1007/s11065-014-9246-9

Gogolla, N. (2017). The insular cortex. Current Biology, 27(12), R580–R586. 10.1016/j.cub.2017.05.010

Goldie, P. (1999). How We Think of Others’ Emotions. Mind & Language, 14(4), 394–423. 10.1111/1468-0017.00118

Gutchess, A., & Kensinger, E. A. (2018). Shared Mechanisms May Support Mnemonic Benefits from Self-Referencing and Emotion. Trends in Cognitive Sciences, 22(8), 712–724. 10.1016/j.tics.2018.05.001

Happé, F., Cook, J. L., & Bird, G. (2017). The Structure of Social Cognition: In(ter)dependence of Sociocognitive Processes. Annual Review of Psychology, 68(1), 243–267. 10.1146/annurev-psych-010416-044046

Henrie, J. A., & Shapley, R. (2005). LFP Power Spectra in V1 Cortex: The Graded Effect of Stimulus Contrast. Journal of Neurophysiology, 94(1), 479–490. 10.1152/jn.00919.2004

Herman, A. B., Smith, E. H., Schevon, C. A., Yates, M. J., McKhann, G. M., Botvinick, M., Hayden, B. Y., & Sheth, S. A. (2023). Pretrial predictors of conflict response efficacy in the human prefrontal cortex. iScience, 26(11), 108047. 10.1016/j.isci.2023.108047

Hultman, R., Mague, S. D., Li, Q., Katz, B. M., Michel, N., Lin, L., Wang, J., David, L. K., Blount, C., Chandy, R., Carlson, D., Ulrich, K., Carin, L., Dunson, D., Kumar, S., Deisseroth, K., Moore, S. D., & Dzirasa, K. (2016). Dysregulation of Prefrontal Cortex-Mediated Slow-Evolving Limbic Dynamics Drives Stress-Induced Emotional Pathology. Neuron, 91(2), 439–452. 10.1016/j.neuron.2016.05.038

Jackson, A. D., Cohen, J. L., Phensy, A. J., Chang, E. F., Dawes, H. E., & Sohal, V. S. (2024). Amygdala-hippocampus somatostatin interneuron beta-synchrony underlies a cross-species biomarker of emotional state. Neuron, 112(7), 1182–1195.e5. 10.1016/j.neuron.2023.12.017

Janak, P. H., & Tye, K. M. (2015). From circuits to behaviour in the amygdala. Nature, 517(7534), Article 7534. 10.1038/nature14188

Joshi, A., Scheinost, D., Okuda, H., Belhachemi, D., Murphy, I., Staib, L. H., & Papademetris, X. (2011). Unified framework for development, deployment and robust testing of neuroimaging algorithms. Neuroinformatics, 9(1), 69–84. 10.1007/s12021-010-9092-8

Kanske, P., Böckler, A., Trautwein, F.-M., & Singer, T. (2015). Dissecting the social brain: Introducing the EmpaToM to reveal distinct neural networks and brain–behavior relations for empathy and Theory of Mind. NeuroImage, 122, 6–19. 10.1016/j.neuroimage.2015.07.082

Karnath, H.-O., Baier, B., & Nägele, T. (2005). Awareness of the Functioning of One’s Own Limbs Mediated by the Insular Cortex? Journal of Neuroscience, 25(31), 7134–7138. 10.1523/JNEUROSCI.1590-05.2005

Koeppel, C. J., Ruser, P., Kitzler, H., Hummel, T., & Croy, I. (2020). Interoceptive accuracy and its impact on neuronal responses to olfactory stimulation in the insular cortex. Human Brain Mapping, 41(11), 2898–2908. 10.1002/hbm.24985

Lachaux, J.-P., Axmacher, N., Mormann, F., Halgren, E., & Crone, N. E. (2012). High-frequency neural activity and human cognition: Past, present and possible future of intracranial EEG research. Progress in Neurobiology, 98(3), 279–301. 10.1016/j.pneurobio.2012.06.008

Lamm, C., Decety, J., & Singer, T. (2011). Meta-analytic evidence for common and distinct neural networks associated with directly experienced pain and empathy for pain. NeuroImage, 54(3), 2492–2502. 10.1016/j.neuroimage.2010.10.014

Luo, Y., Qin, S., Fernández, G., Zhang, Y., Klumpers, F., & Li, H. (2014). Emotion perception and executive control interact in the salience network during emotionally charged working memory processing. Human Brain Mapping, 35(11), 5606–5616. 10.1002/hbm.22573

Magnotti, J. F., Wang, Z., & Beauchamp, M. S. (2020). RAVE: Comprehensive open-source software for reproducible analysis and visualization of intracranial EEG data. NeuroImage, 223, 117341. 10.1016/j.neuroimage.2020.117341

Manning, J. R., Jacobs, J., Fried, I., & Kahana, M. J. (2009). Broadband shifts in local field potential power spectra are correlated with single-neuron spiking in humans. The Journal of Neuroscience: The Official Journal of the Society for Neuroscience, 29(43), 13613–13620. 10.1523/JNEUROSCI.2041-09.2009

Marsh, A. A. (2018). The neuroscience of empathy. Current Opinion in Behavioral Sciences, 19, 110–115. 10.1016/j.cobeha.2017.12.016

Menon, V., & Uddin, L. Q. (2010). Saliency, switching, attention and control: A network model of insula function. Brain Structure & Function, 214(5–6), 655–667. 10.1007/s00429-010-0262-0

Metzger, B. A., Kalva, P., Mocchi, M. M., Cui, B., Adkinson, J. A., Wang, Z., Mathura, R., Kanja, K., Gavvala, J., Krishnan, V., Lin, L., Maheshwari, A., Shofty, B., Magnotti, J. F., Willie, J. T., Sheth, S. A., & Bijanki, K. R. (2023). Intracranial stimulation and EEG feature analysis reveal affective salience network specialization. Brain, awad200. 10.1093/brain/awad200

Miller, K. J., Honey, C. J., Hermes, D., Rao, R. P., denNijs, M., & Ojemann, J. G. (2014). Broadband changes in the cortical surface potential track activation of functionally diverse neuronal populations. NeuroImage, 85, 711–720. 10.1016/j.neuroimage.2013.08.070

Mitchell, R. L. C., & Phillips, L. H. (2015). The overlapping relationship between emotion perception and theory of mind. Neuropsychologia, 70, 1–10. 10.1016/j.neuropsychologia.2015.02.018

Molenberghs, P., Johnson, H., Henry, J. D., & Mattingley, J. B. (2016). Understanding the minds of others: A neuroimaging meta-analysis. Neuroscience & Biobehavioral Reviews, 65, 276–291. 10.1016/j.neubiorev.2016.03.020

Molnar-Szakacs, I., & Uddin, L. Q. (2022). Anterior insula as a gatekeeper of executive control. Neuroscience & Biobehavioral Reviews, 139, 104736. 10.1016/j.neubiorev.2022.104736

Münkler, P., Rothkirch, M., Dalati, Y., Schmack, K., & Sterzer, P. (2015). Biased Recognition of Facial Affect in Patients with Major Depressive Disorder Reflects Clinical State. PLOS ONE, 10(6), e0129863. 10.1371/journal.pone.0129863

Nguyen, V. T., Breakspear, M., Hu, X., & Guo, C. C. (2016). The integration of the internal and external milieu in the insula during dynamic emotional experiences. NeuroImage, 124, 455–463. 10.1016/j.neuroimage.2015.08.078

Nir, Y., Fisch, L., Mukamel, R., Gelbard-Sagiv, H., Arieli, A., Fried, I., & Malach, R. (2007). Coupling between Neuronal Firing Rate, Gamma LFP, and BOLD fMRI Is Related to Interneuronal Correlations. Current Biology, 17(15), 1275–1285. 10.1016/j.cub.2007.06.066

Nowacki, J., Wingenfeld, K., Kaczmarczyk, M., Chae, W. R., Abu-Tir, I., Deuter, C. E., Piber, D., Hellmann-Regen, J., & Otte, C. (2020). Cognitive and emotional empathy after stimulation of brain mineralocorticoid and NMDA receptors in patients with major depression and healthy controls. Neuropsychopharmacology, 45(13), Article 13. 10.1038/s41386-020-0777-x

Paulus, M. P., & Khalsa, S. S. (2021). When You Don’t Feel Right Inside: Homeostatic Dysregulation and the Mid-Insular Cortex in Psychiatric Disorders. American Journal of Psychiatry, 178(8), 683–685. 10.1176/appi.ajp.2021.21060622

Preckel, K., Kanske, P., & Singer, T. (2018). On the interaction of social affect and cognition: Empathy, compassion and theory of mind. Current Opinion in Behavioral Sciences, 19, 1–6. 10.1016/j.cobeha.2017.07.010

Price, R. B., & Duman, R. (2020). Neuroplasticity in cognitive and psychological mechanisms of depression: An integrative model. Molecular Psychiatry, 25(3), Article 3. 10.1038/s41380-019-0615-x

Rajmohan, V., & Mohandas, E. (2007). The limbic system. Indian Journal of Psychiatry, 49(2), 132–139. 10.4103/0019-5545.33264

Rich, E. L., & Wallis, J. D. (2017). Spatiotemporal dynamics of information encoding revealed in orbitofrontal high-gamma. Nature Communications, 8(1), Article 1. 10.1038/s41467-017-01253-5

Rogers-Carter, M. M., Varela, J. A., Gribbons, K. B., Pierce, A. F., McGoey, M. T., Ritchey, M., & Christianson, J. P. (2018). Insular cortex mediates approach and avoidance responses to social affective stimuli. Nature Neuroscience, 21(3), 404–414. 10.1038/s41593-018-0071-y

Rolls, E. T. (2016). Functions of the anterior insula in taste, autonomic, and related functions. Brain and Cognition, 110, 4–19. 10.1016/j.bandc.2015.07.002

Rolls, E. T. (2019). The cingulate cortex and limbic systems for emotion, action, and memory. Brain Structure & Function, 224(9), 3001–3018. 10.1007/s00429-019-01945-2

Samide, R., Cooper, R. A., & Ritchey, M. (2020). A database of news videos for investigating the dynamics of emotion and memory. Behavior Research Methods, 52(4), 1469–1479. 10.3758/s13428-019-01327-w

Schaafsma, S. M., Pfaff, D. W., Spunt, R. P., & Adolphs, R. (2015). Deconstructing and reconstructing theory of mind. Trends in Cognitive Sciences, 19(2), 65–72. 10.1016/j.tics.2014.11.007

Schirmer, A., & Adolphs, R. (2017). Emotion Perception from Face, Voice, and Touch: Comparisons and Convergence. Trends in Cognitive Sciences, 21(3), 216–228. 10.1016/j.tics.2017.01.001

Schurz, M., Radua, J., Tholen, M. G., Maliske, L., Margulies, D. S., Mars, R. B., Sallet, J., & Kanske, P. (2021). Toward a hierarchical model of social cognition: A neuroimaging meta-analysis and integrative review of empathy and theory of mind. Psychological Bulletin, 147(3), 293–327. 10.1037/bul0000303

Segerdahl, A. R., Mezue, M., Okell, T. W., Farrar, J. T., & Tracey, I. (2015). The dorsal posterior insula subserves a fundamental role in human pain. Nature Neuroscience, 18(4), Article 4. 10.1038/nn.3969

Sheth, S. A., Bijanki, K. R., Metzger, B., Allawala, A., Pirtle, V., Adkinson, J. A., Myers, J., Mathura, R. K., Oswalt, D., Tsolaki, E., Xiao, J., Noecker, A., Strutt, A. M., Cohn, J. F., McIntyre, C. C., Mathew, S. J., Borton, D., Goodman, W., & Pouratian, N. (2022). Deep Brain Stimulation for Depression Informed by Intracranial Recordings. Biological Psychiatry, 92(3), 246–251. 10.1016/j.biopsych.2021.11.007

Singer, T., Critchley, H. D., & Preuschoff, K. (2009). A common role of insula in feelings, empathy and uncertainty. Trends in Cognitive Sciences, 13(8), 334–340. 10.1016/j.tics.2009.05.001

Singer, T., & Lamm, C. (2009). The social neuroscience of empathy. Annals of the New York Academy of Sciences, 1156, 81–96. 10.1111/j.1749-6632.2009.04418.x

Singer, T., Seymour, B., O’Doherty, J., Kaube, H., Dolan, R. J., & Frith, C. D. (2004). Empathy for Pain Involves the Affective but not Sensory Components of Pain. Science, 303(5661), 1157–1162. 10.1126/science.1093535

Singer, T., & Tusche, A. (2014). Chapter 27 - Understanding Others: Brain Mechanisms of Theory of Mind and Empathy. In P. W. Glimcher & E. Fehr (Eds.), Neuroeconomics (Second Edition) (pp. 513–532). Academic Press. 10.1016/B978-0-12-416008-8.00027-9

Spunt, R. P., & Adolphs, R. (2019). The neuroscience of understanding the emotions of others. Neuroscience Letters, 693, 44–48. 10.1016/j.neulet.2017.06.018

Surguladze, S. A., Young, A. W., Senior, C., Brébion, G., Travis, M. J., & Phillips, M. L. (2004). Recognition accuracy and response bias to happy and sad facial expressions in patients with major depression. Neuropsychology, 18(2), 212–218. 10.1037/0894-4105.18.2.212

Tottenham, N., Tanaka, J. W., Leon, A. C., McCarry, T., Nurse, M., Hare, T. A., Marcus, D. J., Westerlund, A., Casey, B. J., & Nelson, C. (2009). The NimStim set of facial expressions: Judgments from untrained research participants. Psychiatry Research, 168(3), 242–249. 10.1016/j.psychres.2008.05.006

Tschida, J. E., & Yerys, B. E. (2021). A Systematic Review of the Positive Valence System in Autism Spectrum Disorder. Neuropsychology Review, 31(1), 58–88. 10.1007/s11065-020-09459-z

Uddin, L. Q. (2015). Salience processing and insular cortical function and dysfunction. Nature Reviews Neuroscience, 16(1), Article 1. 10.1038/nrn3857

Uddin, L. Q. (2017). Chapter 2—Anatomy of the Salience Network. In L. Q. Uddin (Ed.), Salience Network of the Human Brain (pp. 5–10). Academic Press. 10.1016/B978-0-12-804593-0.00002-3

Underwood, R., Tolmeijer, E., Wibroe, J., Peters, E., & Mason, L. (2021). Networks underpinning emotion: A systematic review and synthesis of functional and effective connectivity. NeuroImage, 243, 118486. 10.1016/j.neuroimage.2021.118486

Vickery, T. J., Chun, M. M., & Lee, D. (2011). Ubiquity and specificity of reinforcement signals throughout the human brain. Neuron, 72(1), 166–177.

Westlin, C., Theriault, J. E., Katsumi, Y., Nieto-Castanon, A., Kucyi, A., Ruf, S. F., … & Barrett, L. F. (2023). Improving the study of brain-behavior relationships by revisiting basic assumptions. Trends in cognitive sciences, 27(3), 246–257.

Widge, A. S., Heilbronner, S. R., & Hayden, B. Y. (2019). Prefrontal cortex and cognitive control: new insights from human electrophysiology. F1000Research, 8.

Xiao, J., Provenza, N. R., Asfouri, J., Myers, J., Mathura, R. K., Metzger, B., Adkinson, J. A., Allawala, A. B., Pirtle, V., Oswalt, D., Shofty, B., Robinson, M. E., Mathew, S. J., Goodman, W. K., Pouratian, N., Schrater, P. R., Patel, A. B., Tolias, A. S., Bijanki, K. R., … Sheth, S. A. (2023). Decoding Depression Severity From Intracranial Neural Activity. Biological Psychiatry. 10.1016/j.biopsych.2023.01.020

Yoo, S. B. M., & Hayden, B. Y. (2020). The transition from evaluation to selection involves neural subspace reorganization in core reward regions. Neuron, 105(4), 712–724.

Zhao, Y., Zhang, L., Rütgen, M., Sladky, R., & Lamm, C. (2022). Effective connectivity reveals distinctive patterns in response to others’ genuine affective experience of disgust. NeuroImage, 259, 119404. 10.1016/j.neuroimage.2022.119404

Zheng, J., Stevenson, R. F., Mander, B. A., Mnatsakanyan, L., Hsu, F. P. K., Vadera, S., Knight, R. T., Yassa, M. A., & Lin, J. J. (2019). Multiplexing of Theta and Alpha Rhythms in the Amygdala-Hippocampal Circuit Supports Pattern Separation of Emotional Information. Neuron, 102(4), 887–898.e5. 10.1016/j.neuron.2019.03.025

Zhou, F., Zhao, W., Qi, Z., Geng, Y., Yao, S., Kendrick, K. M., Wager, T. D., & Becker, B. (2021). A distributed fMRI-based signature for the subjective experience of fear. Nature Communications, 12(1), Article 1. 10.1038/s41467-021-26977-3

